# Abnormal intrinsic neural timescale and reduced grey matter volume in Parkinson’s disease

**DOI:** 10.1101/2022.09.17.508074

**Authors:** Yarui Wei, Chunyan Zhang, Yuanyuan Peng, Chen Chen, Shaoqiang Han, Weijian Wang, Yong Zhang, Hong Lu, Jingliang Cheng

## Abstract

BACKGROUND: Numerous studies indicate altered peak latency of event-related potential and altered time variability of brain function network in Parkinson’s disease (PD), and the intrinsic neural timescale estimated how long neural information stored in a local brain area has been specialized. However, it was unclear whether PD patients exhibited abnormal intrinsic timescales and accompanied with abnormal grey matter and whether PD patients exhibited different temporal feature at different stages. STUDY TYPE: Prospective. POPULATION: 74 PD patients, including 44 patients in the early stage (PD-ES) and 30 patients in the late stage (PD-LS), and 73 healthy controls (HC). SEQUENCE: 3.0T MRI scanner; prototypic T1 magnetization prepared rapid acquisition gradient echo (MPRAGE); resting-state fMRI. ASSESSMENT: the intrinsic timescales were estimated by using the magnitude of the autocorrelation of intrinsic neural signals. Voxel-based morphometry (VBM) was performed to calculate the grey matter volume (GMV) in the whole brain. STATISTICAL TEST: Analysis of variance (ANOVA); two-sample *t*-tests; Spearman rank correlation analysis; Mann-Whitney U test; Kruskal-Wallis’ H test. RESULTS: We identified that the PD group had abnormal intrinsic timescales in bilateral lingual and calcarine gyri, bilateral postcentral and precentral gyri, and the right middle cingulum gyrus, which correlated with the symptom severity and the GMV. Moreover, longer timescale in the right middle frontal gyrus were also found in the PD group. Increasingly, the PD-ES group had longer timescales in the anterior cortical regions, whereas the PD-LS group had shorter timescales in the posterior cortical regions. DATA CONCLUSION: Our findings suggest that PD patients exhibit abnormal intrinsic timescales in visual, sensorimotor, and cognitive systems and distinct patterns of intrinsic timescales and GMV in cerebral cortex at different stages, which might provide new insights for the neural substrate of PD.

## Introduction

Specialization and hierarchy are organizing principles for primate cortex, and cortical areas are also specialized in the temporal domain. How long neural information stored in a local brain area has been quantified by intrinsic neural timescales. The previous studies estimated the intrinsic neural timescales by using the magnitude of the autocorrelation of intrinsic neural signals^1, 2^. The neural timescale in a local brain area reflects the function of that area in many trials of a task^3, 4^ or even in the absence of direct stimulus processing^5, 6^. Moreover, the intrinsic neural timescales at resting state could predict the neural activity of the brain region in a task^7–9^, and the sensory and prefrontal areas exhibit shorter and longer timescales at resting state, respectively^5^. Recently, the intrinsic timescales were also verified by using multiple neuroimaging methods, such as single-neuron level^7^, electroencephalography (EEG)^10^, magnetoencephalogram (MEG)^9^, and functional magnetic resonance imaging (fMRI)^1, 2, 6, 11^.

Numerous studies also indicate altered temporal features in Parkinson’s disease (PD) using the methods of event-related potentials (ERPs)^12, 13^ and dynamic functional connectivity (DFC)^14–17^. ERPs gained from the scalp-recorded EEG are calculated by averaging EEG activity that is time-locked to the occurrence of an event^18^. As classical ERP components, P3b latency and mismatch negativity (MMN) amplitude are sensitive to PD dementia, and P3a amplitude is a potential marker of PD disease progression^19^. On the other hand, DFC between brain regions assumes spatial and temporal dynamic of fMRI data using sliding window approach^20, 21^. Abnormal time variability of the intrinsic brain functional network was also found in PD^14–17^. Although the upper two neuroimaging methods provide us some evidence related to the temporal feature for neural changes of PD, it’s unclear whether exact intrinsic timescales in local brain regions were abnormal in patients with PD.

PD is a progressive neurological disorder characterized by tremor, rigidity, and slowness of movements and abnormal sensorimotor integration^22, 23^. Nonmotor symptoms, such as visual dysfunction (retinal deficits, facial emotion recognition deficits, and visual hallucinations)^24, 25^ and cognitive impairment^26, 27^, also occur frequently in PD. Resting state fMRI studies also showed abnormal neural activity or functional connection in motor^28–31^ and nonmotor^32–35^ areas. Moreover, studies with somatosensory evoked potentials (SEPs)^36^, prepulse inhibition^37^, and ERPs^38^ support the hypothesis of central abnormalities of sensorimotor integration in PD. Importantly, numerous studies showed that patients with PD with cognitive impairments exhibited changes of brain structure and function^32, 33, 35, 39^. In addition, visual-related areas also showed abnormal neural activation, and structural and functional changes in PD^40–42^. Therefore, we hypothesized abnormal neural information also stored in sensorimotor, visual, and cognitive-related areas in PD patients.

Here, we assessed whether abnormal intrinsic timescales were exhibited in the brain of PD patients. In line with the previous studies^1, 2^, the present study also estimated the intrinsic timescales by using the magnitude of the autocorrelation of intrinsic neural signals in PD patients. Moreover, we also explored the correlation between abnormal intrinsic timescales in PD patients and clinical symptoms. Because the previous studies found significant correlation between abnormal intrinsic timescales and the structural atypicality in autism^1^, and the factor of structural changes of the brain^43–45^ in PD are also important. Therefore, we also explored the correlations between abnormal intrinsic timescales and the structure information in PD patients. Finally, because PD is a progressive neurodegenerative disease and imaging biomarkers in PD are increasingly important for monitoring progression in clinical trials^46, 47^, we also explored the differences of intrinsic timescales among patients with PD in the early stage (PD-ES) and the late stage (PD-LS) and healthy controls (HC).

## Materials and Methods

The present research was approved by the Ethics Committee of our institution. All subjects or their guardians signed the informed consent forms after the nature of the procedures had been fully explained.

### Participants

A total of 74 PD patients (41 females) and 73 age- and sex-matched HC (35 females) were included in the study (Table 1). The clinical diagnosis of patients with PD was made by three experienced neurologists with more than five years of experience according to the clinical diagnostic criteria for PD introduced by the International Movement Disorder Society (MDS)^48^, and all patients were diagnosed as idiopathic PD. PD patients were assessed the severity of motor symptoms and cognitive impairments by using the unified Parkinson’s disease rating scale (UPDRS), a revised severity classification Hoehn and Yahr (HY) scale, Montreal cognitive assessment (MoCA), and mini-mental state examination (MMSE). Based on HY scale, 74 patients with PD were split into 44 patients with PD-ES (the scores of the HY scale were ≤2) and 30 patients with PD-LS (the scores of the HY scale were >2) (Supplementary table 1). Moreover, we collected UPDRS data for 72 of the patients, and collected HY scale data for all patients, and collected MoCA data for 57 of the patients, and collected MMSE data for 71 of the patients. The scores of MoCA and MMSE lower than 25 considered cognitive impairment^49^. Exclusion criteria for the patients were as follows: (1) secondary or atypical parkinsonism; (2) history of other neurological diseases (cerebrovascular disease, seizures, brain surgery, brain tumors, moderate to severe head trauma, or hydrocephalus); (3) history of psychiatric illness; (4) use of antipsychotics or other medicines that may affect clinical assessment; (5) alcohol or drug abuse; (6) contraindications for MRI. The HC were healthy middle-aged and elderly people recruited from across Henan Province. Exclusion criteria for HC were as follows: (1) history of neurological or psychiatric illness; (2) alcohol or drug abuse; (3) contraindications for MRI. Because we performed the Shapiro-Wilk test to check the normality of demographic and clinical data for each group, and the results showed that the factor of demographic and clinical data was not normally distributed for each group. Therefore, Wilcoxon signed-rank test were used to compare demographic and clinical data between groups and *X*^2^ test were used to compare sex data between groups with SPSS software, respectively.

### Data acquisition

All subjects were scanned using one of two 3T MAGNETOM Prisma scanners (Siemens Healthcare, Erlangen, Germany), with a 64-channel receiver array head coil. Head motion and scanner noise were reduced using foam paddings and earplugs, respectively. All participants were asked to remain alert, not to move, not to sleep with eyes closed, not to think systematically. All patients have been scanned during the ON phase. Although scanning during the OFF phase might increase the disease effect it makes it harder to scan due to motion artifacts. Therefore, we preferred to scan during the ON phase to decrease motion artifacts. Structural images were acquired using a 3D prototypic T1 magnetization prepared rapid acquisition gradient echo (MPRAGE) sequence with the following settings: repetition time (TR)/echo time (TE) = 2300/2.9 ms, slice number = 176, slice thickness = 1.2 mm, slice gap = 0 mm, flip angle = 9°, field of view (FOV) = 24.0 × 25.6 cm^2^, number of averages = 1, matrix size = 240 × 256, voxel size = 1 × 1 × 1.2 mm^3^, scan time = 5.2 mins. Functional images were acquired transversely with echo planar imaging (EPI) sequence with the following settings: TR/TE = 1000/30 ms, slice number = 52, slice thickness = 2.2 mm, slice gap = 2.64 mm, flip angle = 70°, FOV = 22 × 22 cm^2^, number of averages = 1, matrix size = 64 × 64, voxel size = 3.4375 × 3.4375 × 2.2 mm^3^. A total of 400 volumes were collected, resulting in a total scan time of 6.67 mins.

### Data preprocessing

The functional images were preprocessed using Data Processing Assistant for Resting-State fMRI (DPARSF) programs, which are based on Statistical Parametric Mapping (SPM12, http://www.fil.ion.ucl.ac.uk/spm) and MATLAB (MathWorks). The first 5 volumes were discarded due to unsteady magnetization. Slice-timing and realignment were performed. Then, data were spatially normalized to the Montreal Neurological Institute (MNI) template (resampling voxel size = 3 × 3 × 3 mm^3^), detrended, and filtered (0.01-0.08 Hz). Image volumes with framewise displacement (FD) >0.2 mm were scrubbed to reduce the effect of head motion using spline interpolation. Because we performed the Shapiro-Wilk test to check the normality of FD for each group and found that FD was not normally distributed for each group. No significant between-group difference in FD was found for the PD and HC groups (*P* = 0.109) and for the PD-ES and PD-LS groups (*P* = 0.991).

Grey matter volume (GMV) was calculated from structural MRI data using SPM12 and DPABI as follows: firstly, we segmented the structural images into grey matter, white matter, and cerebrospinal fluid; then, segmented images were also spatially normalized to the MNI template (resampling voxel size = 2 × 2 × 2 mm^3^) and modulated; finally, the modulated gray matter maps were smoothed using an isotropic Gaussian kernel [full width at half maximum (FWHM) = 6 mm].

### Intrinsic timescale map

Based on the previous study^1^, we used the preprocessed fMRI data to evaluate the intrinsic neural timescale for each voxel in the whole brain of each participant as follows. First, we estimated an autocorrelation function (ACF) of the fMRI signal of each voxel (time bin = TR), and then calculated the sum of ACF values in the initial period where the ACF showed positive values. The upper limit of this period was set at the point where the ACF hits zero for the first time. After repeating this procedure for every voxel, we also spatially smoothed the brain map (FWHM = 6 mm) to improve the signal-to-noise ratio and then obtained *Z*-transformed brain map to eliminate individual differences within group. We used this whole-brain map as an intrinsic timescale map in which the value at each voxel is equal to the intrinsic neural timescale of the brain region. Two-sample *t* test was performed to compare the group differences in intrinsic timescales between the PD and HC groups, with age, sex, and mean FD as covariates. One-way ANOVA was also performed to compare the group differences in intrinsic timescales between the PD-ES, PD-LS, and HC groups, with age, sex, and mean FD as covariates. Then, post hoc comparisons using Bonferroni’s test were performed, and pair-wise different *Z*-value maps corresponding to between-group intrinsic timescale differences were computed. The statistically significant threshold was set at voxel-wise *P* < 0.001, cluster-wise *P* < 0.05, two-tailed, and the minimum cluster size of 22 voxels after Gaussian random field (GRF) correction.

### Correlations between intrinsic timescale and symptom severity

To investigate the correlations between abnormal intrinsic timescale and symptom severity, we extracted the mean intrinsic timescale values of all voxels within each cluster from corrected statistical map and calculated the Spearman rank correlation coefficients for clinical measures (UPDRS, HY, MoCA, and MMSE) with significant results between groups. Multiple comparisons were corrected using the Bonferroni method (*P* < 0.05/70 = 0.0007).

### Grey matter volume and intrinsic timescale

We designed to investigate the neuroanatomical bases for intrinsic timescale by comparing GMV to the temporal property of neural signals. Two-sample *t* test was performed to compare the group differences in GMV between the PD and HC groups, and one-way ANOVA was also performed to compare the group differences in intrinsic timescales between the PD-ES, PD-LS, and HC groups, with age and sex as covariates. The statistically significant threshold was set at voxel-wise *P* < 0.001, cluster-wise *P* < 0.05, two-tailed, and the minimum cluster size of 58 voxels after GRF correction. We also extracted the mean GMV value of all voxels within each significant cluster from corrected statistical intrinsic timescale maps, performed Mann-Whitney U test to compare the group differences of GMV between the PD and HC groups, and performed Kruskal-Wallis’ H test with Dunn’s test as the post hoc test compare the group differences of GMV among the PD-ES, PD-LS, and HC groups. To investigate the correlations between abnormal intrinsic timescale and GMV, we also extracted the mean intrinsic timescale values of all voxels within each cluster from corrected statistical map and calculated the Spearman rank correlation coefficients for GMV with significant results between groups. Multiple comparisons were corrected using the Bonferroni method (*P* < 0.05/70 = 0.0007).

## Results

### Demographic and clinical data

For the PD and HC groups, no significant between-group difference in age (*P* = 0.810) or sex (*P* = 0.365) was found (Table 1). For the PD-ES and PD-LS groups, significant between-group differences in age, sex, and the scores of UPDRS, HY scale, MoCA and MMSE were found (Supplementary table 1).

### Abnormal intrinsic timescales between the PD and HC groups

The mean intrinsic timescale maps in the PD and HC groups were presented in Fig. 1. Both PD and HC groups had a similar whole-brain pattern of intrinsic timescales: longer timescales in frontal and parietal cortices and shorter timescales in sensorimotor, visual, and auditory areas (Fig. 1, *upper*).

**Figure 1.**
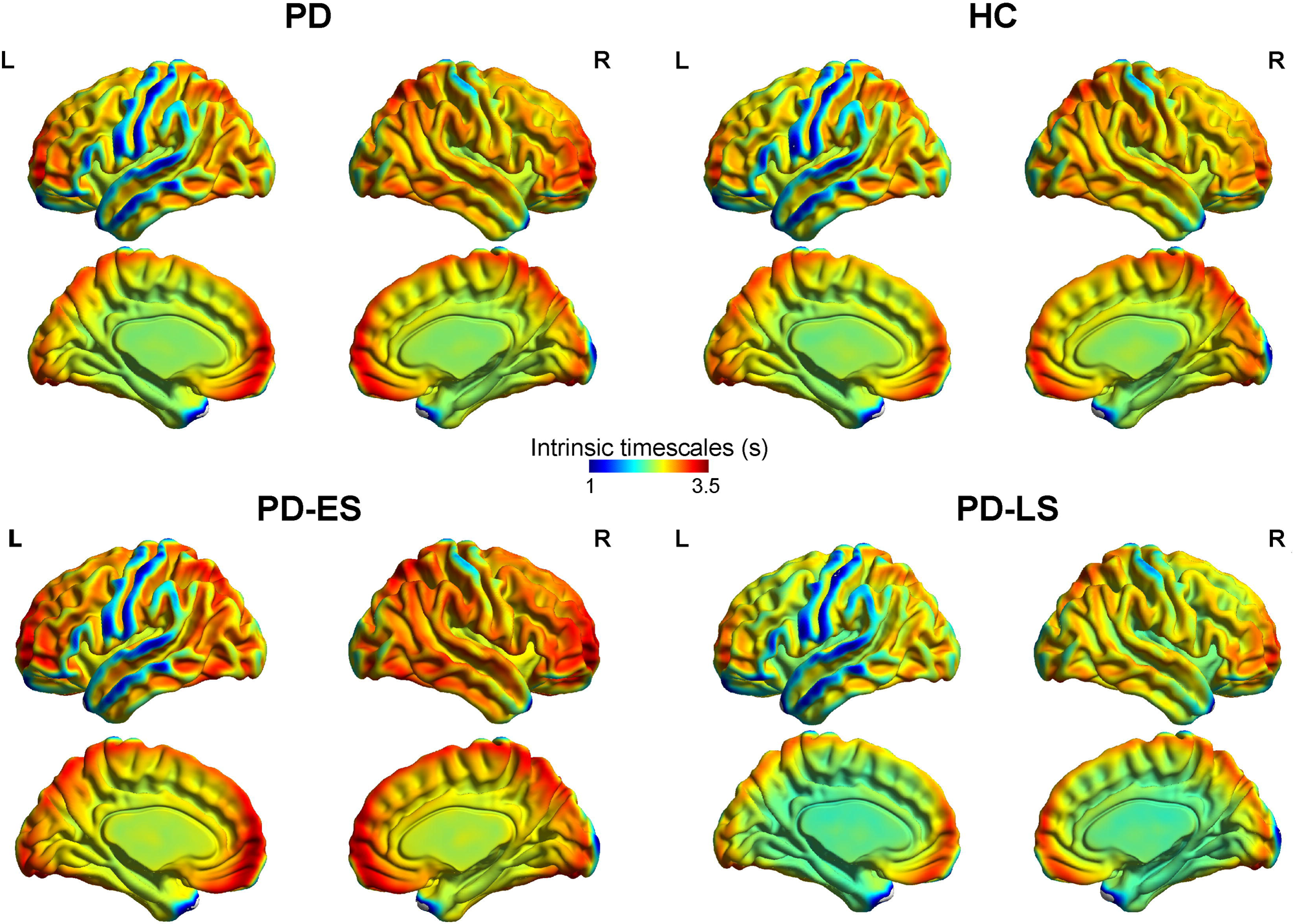
The intrinsic timescales of the whole brain in all PD patients and the HC group, and in the PD patients at different stages. In each group, we found longer intrinsic timescales in frontopartial regions and shorter intrinsic timescales in sensorimotor, visual, and auditory regions. ES, early stage; HC, healthy controls; L, left; LS, late stage; PD, Parkinson’s disease; R, right.

PD group had shorter timescales in the bilateral lingual gyri, bilateral postcentral gyri, and right middle cingulum gyrus compared with the HC group (Supplementary table 2, Fig. 2). The upper first two clusters also included bilateral calcarine gyri and bilateral precentral gyri. Longer timescale in the right middle frontal gyrus was also found in the PD group than the HC group (Supplementary table 2, Fig. 2).

**Figure 2.**
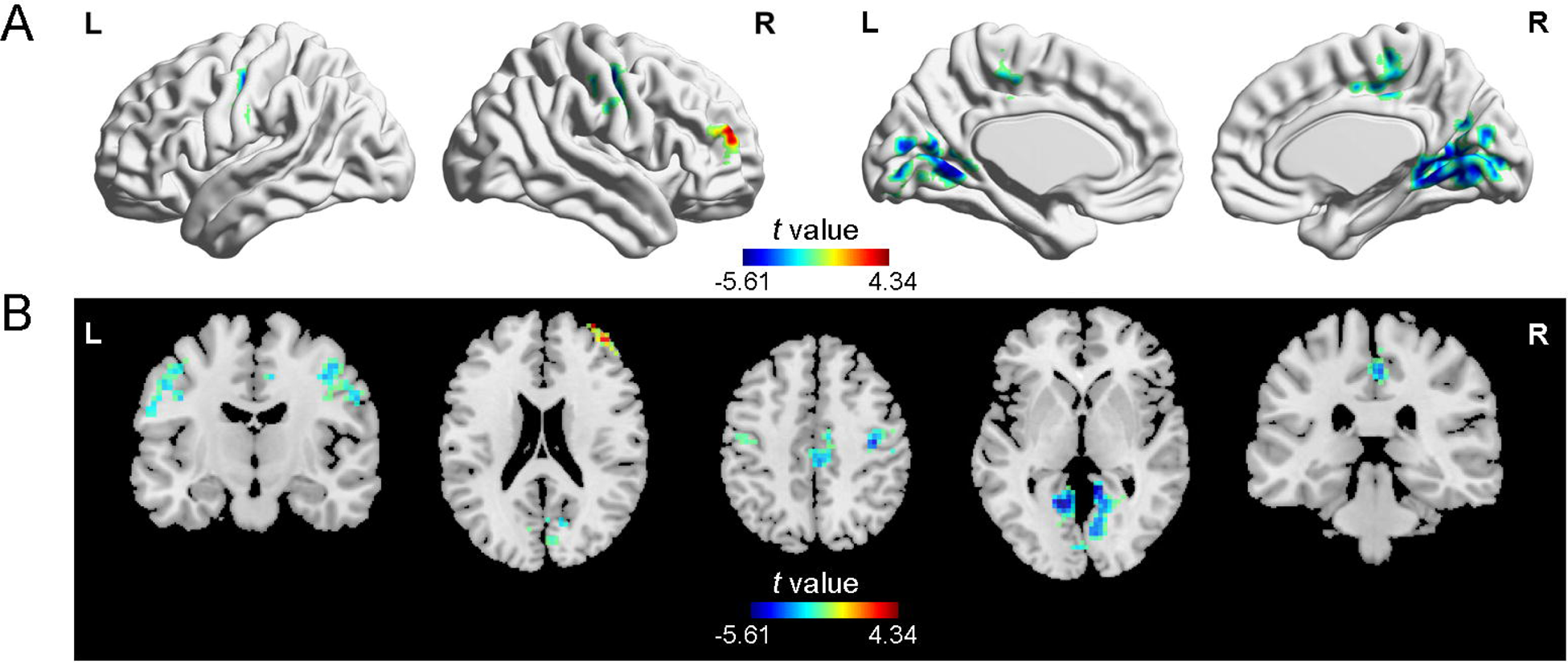
Voxel-wise intrinsic timescales between the PD and HC groups. The PD group had shorter timescales in bilateral postcentral and precentral gyri, bilateral lingual and calcarine gyri, and the right middle cingulum gyrus, and longer timescales in the right middle frontal gyrus than the HC group. HC, healthy controls; L, left; PD, Parkinson’s disease; R, right.

### Correlations between intrinsic timescale and symptom severity in the PD group

The intrinsic timescale of the bilateral lingual gyri in the PD group negatively correlated with the scores of the HY scale (*P* = 0.022, Fig. 3). However, the upper significant results between intrinsic timescale and symptom severity did not remain after Bonferroni correction (*P* > 0.05/70 = 0.0007). No other significant correlation was found between intrinsic timescale and symptom severity in the PD group.

**Figure 3.**
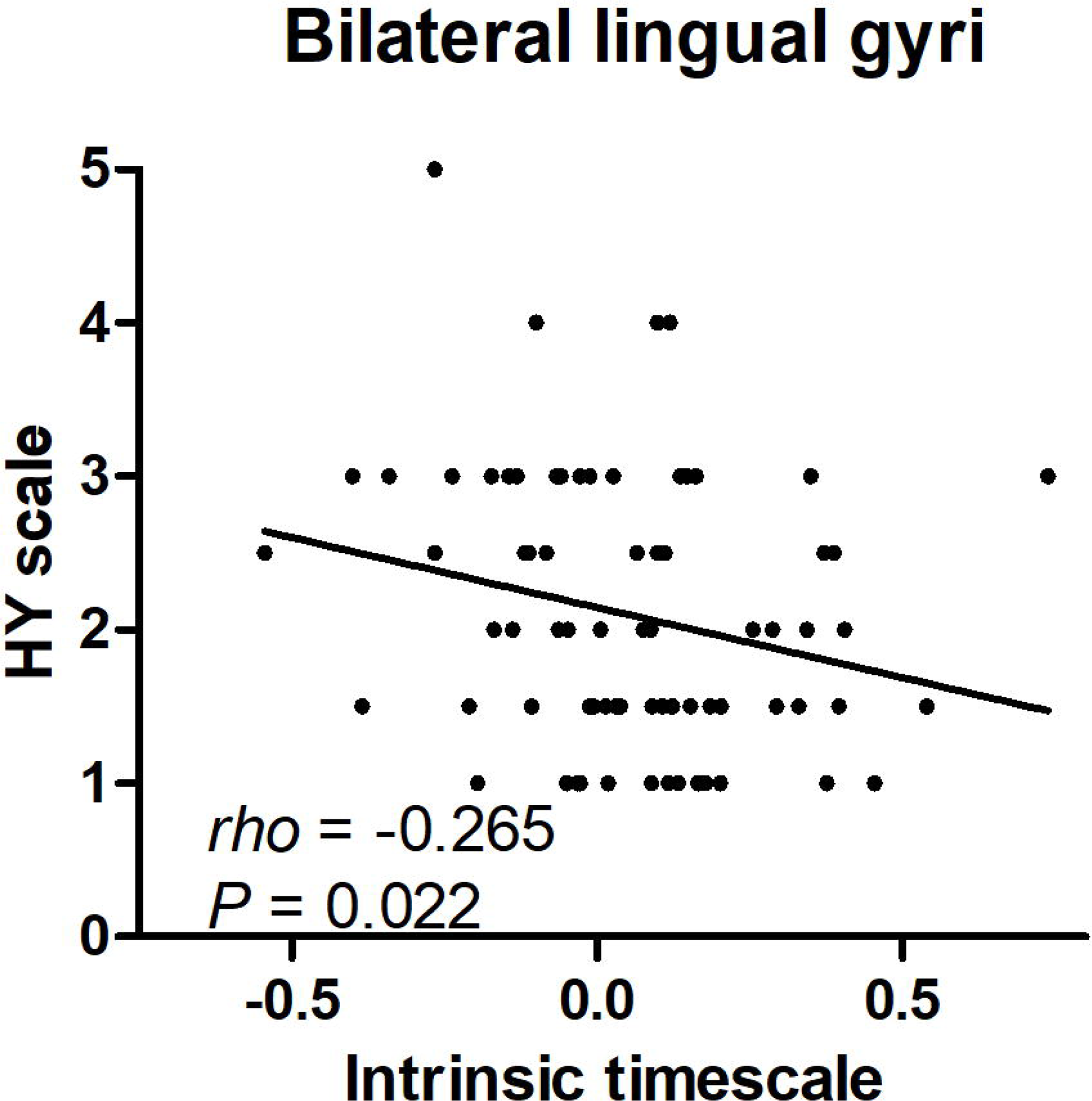
Correlation between the intrinsic timescale and the symptom severity of PD. We found negative correlation between the intrinsic timescale of the lingual gyrus and the scores of the Hoehn and Yahr (HY) scale in patients with Parkinson’s disease (PD).

### Grey matter volume and intrinsic timescale in the PD and HC groups

No significant difference of GMV was found between the PD and HC groups. However, we extracted the mean GMV values of all voxels within each cluster from corrected statistical intrinsic timescale map and found positively significant correlation between the intrinsic timescale and the GMV in the right postcentral gyrus (Fig. 4). However, this significance did not remain after Bonferroni correction (*P* > 0.05/70 = 0.0007). No other significant correlation was found between intrinsic timescale and GMV in the PD group.

**Figure 4.**
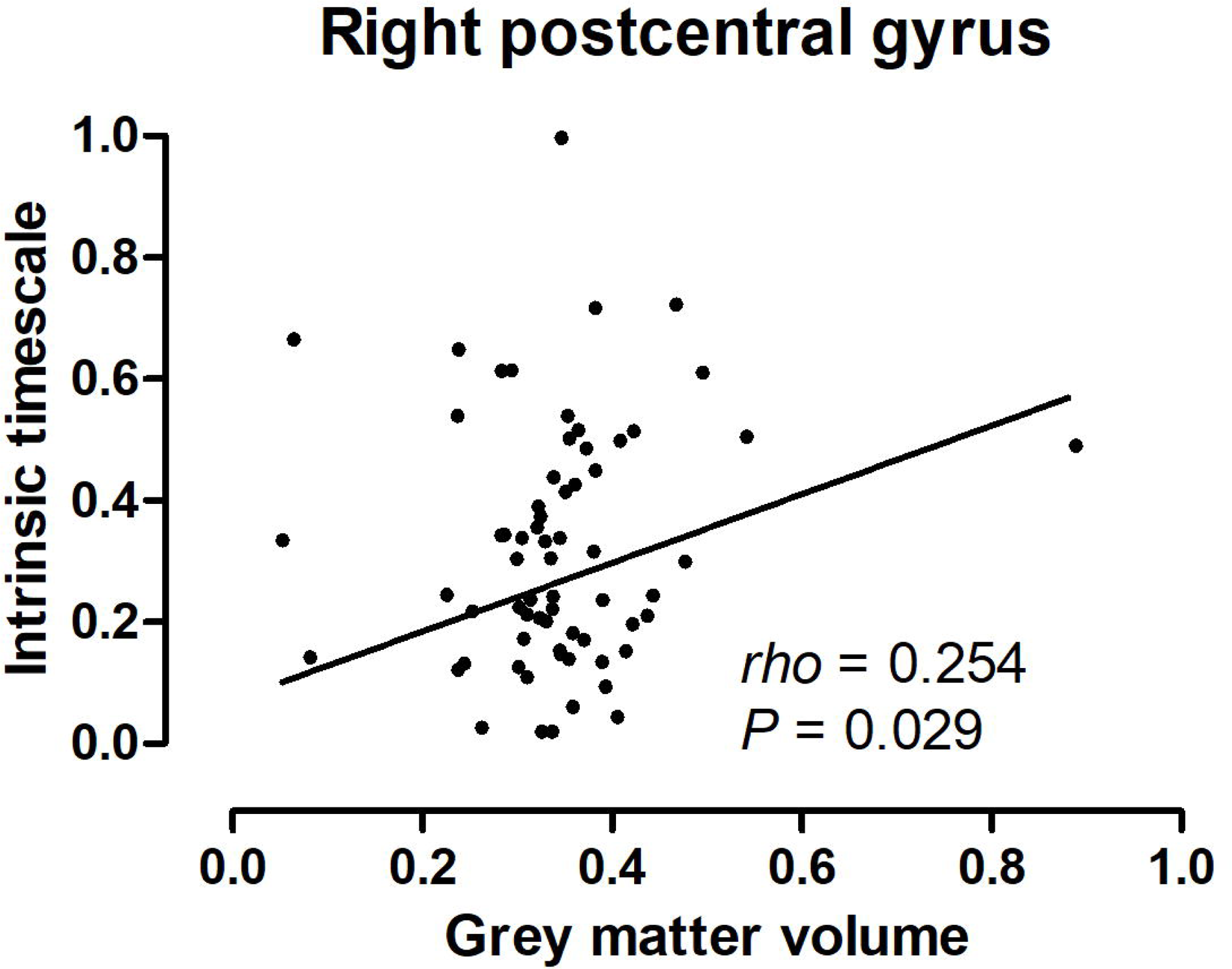
Correlations between the intrinsic timescale and the grey matter volume. Shorter intrinsic timescale in the right postcentral gyrus correlated with its reduced grey matter volume.

### Abnormal intrinsic timescales in PD at different stages

The mean intrinsic timescale maps for the PD-ES and PD-LS groups were also presented in Fig. 1. Both PD-ES and PD-LS groups also showed a similar whole-brain pattern of intrinsic timescales like the PD and HC groups (Fig. 1, *lower*).

The one-sample ANOVA revealed significant between-group differences in intrinsic timescales of bilateral lingual gyri, the left precuneus, and the right middle cingulum gyrus (Supplementary table 3, Fig. 5). The upper first two clusters also included bilateral calcarine gyri. Post hoc comparisons using Bonferroni’s test showed that the PD-LS group had shorter timescale in the left precuneus than that in the PD-ES group and shorter timescales in bilateral calcarine and lingual gyri than that in the HC group (Supplementary table 3, Fig. 5). Increasingly, longer timescales in the left superior frontal gyrus, the left inferior frontal gyrus, the right middle frontal gyrus were also found in the PD-ES group than the HC group (Supplementary table 3, Fig. 5). However, no significant correlation was found between the intrinsic timescale and symptom severity in the PD-ES or PD-LS group.

**Figure 5.**
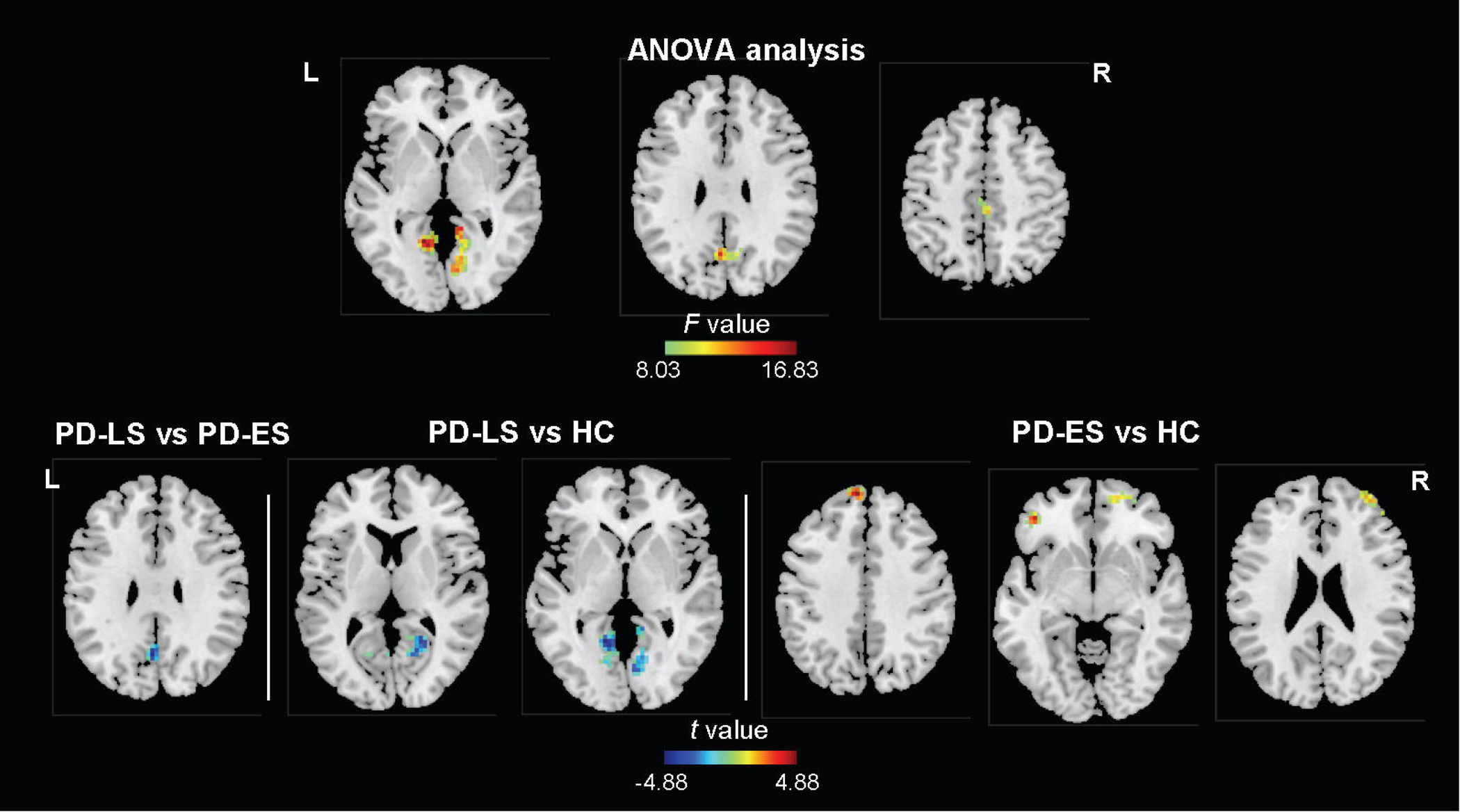
Voxel-wise intrinsic timescales in PD at different stages. One-sample analysis of variance (ANOVA) revealed significant between-group differences in intrinsic timescales of bilateral lingual and calcarine gyri, the left precuneus, and the right middle cingulum gyrus (*Upper*). Post hoc comparisons using Bonferroni’s test showed that the PD-LS group had shorter timescale in the left precuneus than the PD-ES group and shorter timescales in bilateral lingual and calcarine gyri than that in the HC group (*Lower*), and that the PD-ES group had longer timescales in the left superior frontal gyrus, the left inferior frontal gyrus, and the right middle frontal gyrus than the HC group (*Lower*). ES, early stage; HC, healthy controls; L, left; LS, late stage; PD, Parkinson’s disease; R, right

### Grey matter volume and intrinsic timescale in PD at different stages

No significant difference of GMV was found among the PD-ES, PD-LS, and HC groups. However, we found significant differences of mean GMV values between groups in all the upper regions (Table 2). Then we found that: (1) compared to the PD-ES and HC groups, the PD-LS group had reduced GMV in the bilateral lingual and calcarine gyri, and the left precuneus (Table 2). Moreover, compared to the PD-ES group, the PD-LS group also had reduced GMV in the left superior frontal gyrus, the left inferior frontal gyrus, the right middle frontal gyrus and higher GMV in the right middle cingulum gyrus (Table 2). However, no significant correlation was found between the intrinsic timescale and the GMV in the PD-ES or PD-LS group.

## Discussion

In this study, we explored the intrinsic timescales, which relates to the functional hierarchy of the brain^5, 50^, in PD patients. By analyzing the time-dependent magnitude of the autocorrelation function in resting state fMRI data, our findings indicate that PD patients exhibit abnormal intrinsic timescales in motor and non-motor regions, and at different stages, distinct patterns of intrinsic timescales in cerebral cortex, which support the model of motor and non-motor circuit disturbances in PD^51^. The recent study indicates that motor dysfunction of PD is represented as antagonistic interactions within multi-level brain systems^52^.

Our findings also showed abnormal intrinsic timescales in visual, sensorimotor, and cognitive systems, which was also consistent with the studies using the method of DFC. For example, the previous fMRI studies that calculated the variability of brain signal across time found abnormal variance in visual^53–55^, sensorimotor^53, 56^, and frontal^17, 57^ regions in PD. Some studies also found that the symptom severity of PD significantly correlated with such abnormal variability in the visual and sensorimotor regions^55, 56^. Moreover, shorter or longer timescales indicate the time of neural information stored in some regions was shorter and longer, which might explain longer and shorter P3b latency^12, 58, 59^. We speculate that if abnormal signal variability in these regions indicates more random activity of the brain and then result in weaker autocorrelation, which can be interpreted the current findings. Moreover, abnormal intrinsic timescales in multisystem also refer to abnormal functional hierarchy of the brain in individuals with PD. Abnormal intrinsic timescales were also found in some neurological or mental diseases, such as schizophrenia^60, 61^, autism^1^, epilepsy^62^, and depression^63^, which showed that intrinsic timescales might bring new insights about abnormal local neural dynamics on the brain of patients with some diseases. Therefore, the intrinsic timescale might be also a suitable index to explore the neural substrate of PD.

Furthermore, using the index of the intrinsic timescale, we also found that distinct patterns of intrinsic timescales in cerebral cortex in PD at different stages. The cognitive impairment of the PD-LS group was more severity than that of the PD-ES group. Distinct patterns of local cerebral glucose metabolism were also found in PD with and without mild cognitive impairment (PDNC and PDMCI): the PDNC patients had limited areas of hypometabolism in the frontal and occipital cortices, and the PDMCI patients had extensive areas of hypometabolism in the posterior cortical regions, including the temporo-parieto-occipital junction, medial parietal, and inferior temporal cortices^64^. Hirano et al.^65^ suggested that the frontal cortex was associated with impulse control disorders and that posterior brain areas were related to cognitive decline in PD. Distinct patterns of intrinsic timescales between the PD-ES and PD-LS groups also indicate that intrinsic timescale might be a new neuroimaging index across disease stage in PD, which might have the potential to improve clinical care and management.

The GMV and intrinsic timescales might be independent and complementary measures for PD. On the one hand, reduced GMV of bilateral lingual and calcarine gyri, the left precuneus, the right middle cingulate gyrus, and bilateral frontal gyri were found in the PD-LS group, but not in the PD-ES group or in the combined group including all patients at different stages, which could explain the result of no significant difference of GMV between the PD and HC groups. However, the PD-LS group had shorter intrinsic timescales in bilateral lingual and calcarine gyri, and the left precuneus, and the PD-ES group had higher intrinsic timescales in bilateral frontal gyri. Therefore, changes of the GMV in the upper brain regions were consistent and stable for PD patients, and changes of intrinsic timescales were different for PD patients, which might indicate abnormal functional hierarchy in PD. On the other hand, shorter intrinsic timescale in the lingual gyrus correlated with reduced GMV in PD patients, and the latter might be the structure base of abnormal temporal pattern, which was consistent with the previous study^1, 66, 67^. Therefore, a combination of the GMV and intrinsic timescale approaches may provide more evidence for the neural substrate of PD.

Several limitations should be noted when interpreting our findings. First, although we totally recruited 74 PD patients and 73 HC, the sample size is small. Maybe this is the reason why no significant correlation between abnormal intrinsic timescales and symptom/GMV was found in PD at different stages. Abnormal intrinsic timescales in the sensorimotor regions were not found in PD at different stages, which might also originate from the small sample size. The future study should increase the sample size to make the findings more reliable. Second, since abnormal intrinsic timescales of visual regions were obvious in PD, especially in the PD-LS group, detailed information about vision of the participants should have been collected to ensure whether temporal feature in visual regions associated with visual dysfunction. Abnormal neural activity of the visual regions and the symptoms of visual hallucination and other visual dysfunction were also found in PD^40, 68^. Third, significant difference of age and sex was found between the PD-ES and PD-LS groups. However, PD patients were not always ready for recruitment, and their availability would have been reduced even further if the factor of the sex had been applied. The further study should match the factor of the sex between the PD-ES and PD-LS groups, which makes the findings more reliable.

## Conclusions

Our findings suggest that PD patients exhibit abnormal intrinsic timescales in visual, sensorimotor, and cognitive systems and distinct patterns of intrinsic timescales in cerebral cortex in PD patients at early and late stages, which might provide new insights for the neural substrate of PD.

## Data availability statement

The data that support the findings of this study are available from the corresponding author upon reasonable request.

